# Quantification of telomere features in tumor tissue sections by an automated 3D imaging-based workflow

**DOI:** 10.1101/053132

**Authors:** Manuel Gunkel, Inn Chung, Stefan Wörz, Katharina I. Deeg, Ronald Simon, Guido Sauter, David T.W. Jones, Andrey Korshunov, Karl Rohr, Holger Erfle, Karsten Rippe

## Abstract

The microscopic analysis of telomere features provides a wealth of information on the mechanism by which tumor cells maintain their unlimited proliferative potential. Accordingly, the analysis of telomeres in tissue sections of patient tumor samples provides can be exploited to obtain diagnostic information and to define tumor subgroups. In many instances, however, analysis of the image data is conducted by manual inspection of 2D images at relatively low resolution for only a small part of the sample. As the telomere feature signal distribution is frequently heterogeneous, this approach is prone to a biased selection of the information present in the image and lacks subcellular details. Here we address these issues by using an automated high-resolution imaging and analysis workflow that quantifies individual telomere features on tissue sections for a large number of cells. The approach is particularly suited to assess telomere heterogeneity and low abundant cellular sub-populations with distinct telomere characteristics in a reproducible manner. It comprises the integration of multi-color fluorescence in situ hybridization, immunofluorescence and DNA staining with targeted automated 3D fluorescence microscopy and image analysis. We apply our method to telomeres in glioblastoma and prostate cancer samples, and describe how the imaging data can be used to derive statistically reliable information on telomere length distribution or colocalization with PML nuclear bodies. We anticipate that relating this approach to clinical outcome data will prove to be valuable for pretherapeutic patient stratification.

**Abbreviations:**

3D-TIM: 3D targeted imaging
ALT: alternative lengthening of telomeres
APB: ALT-associated PML-NB
CLSM: confocal laser scanning fluorescence microscopy
ECTR: extrachromosomal telomeric repeat
FFPE: formalin-fixed, paraffin-embedded
FISH: fluorescence in situ hybridization
IF: Immunofluorescence
pedGBM: pediatric glioblastoma
PML: promyelocytic leukemia
PML-NB: PML nuclear body
PNA: peptide nucleic acid
ROI: region of interest
TMA: tissue microarray
TMM: telomere maintenance mechanism
SMLM: single molecule localization microscopy

## 1. Introduction

Telomeres are the ends of linear chromosomes and in humans consist of the repetitive sequence 5’-TTAGGG-3’ that is bound by the shelterin protein complex [1]. These repeats are critical for cells, since they prevent the DNA damage signaling machinery from recognizing chromosome ends as double-strand breaks. With every round of replication, however, the number of telomeric repeats decreases. Once the telomere length reaches a critical limit the cells either enter a state called replicative senescence or go into apoptosis. Hence, the continuous shortening of telomeres presents an effective control mechanism to restrict a cell’s potential to divide. In order to proliferate indefinitely, cancer cells have to circumvent this control point by acquiring a telomere maintenance mechanism (TMM) to extend their telomeres. In most tumors, the enzyme telomerase that catalyzes the *de novo* synthesis of telomeric sequences and is silenced in somatic cells is reactivated. In addition to the deregulation of several hundred genes that have been associated with aberrant telomerase activity, de-repression of telomerase can also be caused by mutations in the promoter of the telomerase gene *TERT* as well as structural rearrangements of *TERT* enhancers [2–4]. Apart from the various telomerase related pathways in cancer, alternative lengthening of telomeres (ALT) mechanisms exist in 25-60% of sarcomas, 5-15% of carcinomas and ~25% of glioblastomas (GBMs) [5–7]. These operate in the absence of telomerase and are based on DNA recombination and repair processes. ALT has repressed telomerase gene expression as an overarching characteristic feature and is typically associated with (i) heterogeneous telomere length distribution within individual cells and across tumor cell populations [8], (ii) formation of ALT-associated promyelocytic leukemia (PML) nuclear bodies (APBs) [9, 10], (iii) mutations in genes like *ATRX* and *TP53* as shown for cell lines [11] and through deep-sequencing studies in GBMs, pancreatic neuroendocrine tumors and in other cancers [5, 12, 13], (iv) extrachromosomal telomeric repeats (ECTRs) [14, 15], (v) variant telomeric repeat sequences [16] as well as (vi) high levels of the non-coding telomeric RNA transcript TERRA [17, 18].

Notably, the TMM type can define distinct tumor subgroups that have been linked to patient subgroups and clinical parameters in several entities [3, 4, 19–22]. Thus, understanding telomere maintenance networks provides highly valuable and unique information about the cancer disease state that can be exploited for patient stratification and targeted therapies. This creates the need for methods to identify telomere features and TMMs from primary tumor samples. Often such samples are only available as formalin-fixed, paraffin-embedded (FFPE) tissue sections. These sections are collected either as individual samples or in the form of tissue microarrays (TMAs), where several hundred patient samples are combined on a single microscopy slide. The main method for detection of telomeres on these tissue sections is fluorescence in situ hybridization (FISH) with peptide nucleic acid (PNA) probes that recognize the telomeric repeat sequence 5’-TTAGGG-3’ [23–25]. Once these probes reach the inside of the nucleus they hybridize with the complementary DNA strand with high affinity. The intensity of the fluorescently labeled telomere probe is proportional to the telomere repeat length. Frequently, telomere FISH analysis is used to identify ALT positive tumors. This application is based on the observation that so-called ultra-bright telomere foci of sizes in the range of several µm show a high correlation with the presence of ALT [5, 26]. To this end, telomere FISH-stained tumor sections are manually inspected for the occurrence of FISH signals that are abnormally large and bright. However, it should be noted that the biological source of these ultra-bright foci is unclear. In addition, the current standard analysis uses 2D wide-field microscopy images and lacks structural information along the optical z-axis. Accordingly, tumor sections are classified only in a qualitative or semi-quantitative manner such as ‘short’ and ‘long’ telomeres. Such semi-quantitative analyses have been used to identify correlations between telomere length, tumor reoccurrences and patient survival in several cancer types including prostate cancer, breast cancer and neuroblastoma [24, 27–29].

In our previous work we have introduced a platform for 3D imaging-based quantitative analysis of telomere features by confocal laser scanning fluorescence microscopy (CLSM) [30, 31]. In these two studies an automated three-color confocal RNAi screening platform was developed that allowed us to analyze telomeres and colocalizations between telomeres and PML nuclear bodies (PML-NBs) in cell lines to investigate the ALT mechanism. Based on image analysis the effect of RNAi-mediated knockdown of specific genes on the number and size of telomeres, PML-NBs, and APBs for each individual cell was determined. These features were also related to the cell cycle state as determined by DNA content measured using DAPI staining. Both the image acquisition and data analysis evaluated telomere features in three dimensions. As an extension of this previous work, we here describe an integrated approach to quantify these telomere features on tumor tissue sections and TMAs and illustrate its application to pediatric glioblastoma (pedGBM) and prostate cancer samples. The workflow comprises three main parts: (i) An optimized protocol for fluorescence labeling of tissue sections. (ii) A combination of 2D imaging with a wide-field setup and high-resolution multicolor 3D confocal fluorescence microscopy termed 3D-TIM for 3D Targeted IMaging. An additional zoom-in step by single molecule localization super-resolution microscopy can be included that provides an additional ~10-fold improvement of the resolution. (iii) Quantitative high-resolution 3D image analysis of telomere features to yield statistically reliable measurements for primary tumor samples. The resulting data can be related to clinical data and provide information for pretherapeutic patient stratification.

## 2. Material and Methods

### 2.1. Preparation of tissue sections

In the present work two types of samples were used, namely tissue sections from pediatric glioblastoma samples and prostate cancer TMAs. FFPE sections (5 µm thickness) were prepared in the standard manner from pediatric glioblastoma tissue blocks. Tissues were assessed for tumor cellularity prior to selection, to ensure the presence of >70% tumor cells. The TMA manufacturing process for the prostate cancer samples studied here was as described in detail previously [32].

### 2.2. PNA FISH on prostate cancer TMA

Tissue microarrays were deparaffinized by incubating them three times in xylene for 10 min, incubated twice in 96% ethanol for 5 min and dried at 48°C for 3 min. Next, TMAs were treated with 1 mg/ml proteinase K in TBS for 4 h at 37 °C, washed twice with H_2_O for 3 min, shortly immersed in 96 % ethanol and air dried for a few minutes. Denaturation and FISH probe hybridization was performed as follows: PNA-Hyb solution (70 % formamide, 10 mM Tris HCl, pH 7.5, 0.1 µg/ml salmon sperm) containing 0.1 µM of a Cy3-labeled telomere probe (CCCTAA)_3_ (TelC-Cy3, Panagene) and of a FAM-labeled PNA probe (ATTCGTTGGAAACGGGA) that is directed against the CENP-B binding site in the centromeric alpha satellite DNA (CENP-B-FAM, PNA Bio) was added to the TMA, the slide was heated to 80 °C for 5 min for denaturation and hybridization took place at RT in a wet chamber. Next, the TMAs were washed twice for 15 min in PNA wash buffer (70 % formamide, 10 mM Tris-HCl, pH 7.5), 1 min in 2x SSC, 5 min in 0.1x SSC at 55 °C, 2 x 5 min in 0.05 % Tween-20/ 2x SSC and 3 times in PBS for 5 min. After incubation in 70 %, 85 %, and 100 % ethanol, the slide was air dried and mounted with Prolong reagent including DAPI.

### 2.3. PNA FISH and immunofluorescence on FFPE pedGBM tissue sections

Paraffin was removed from the tissue slices of 5 µm thickness by melting at 65 °C and washing the slides in Xylene. After hydration through a grade ethanol series, slides were incubated in 1 % Tween-20 for 1 min before antigen masking, for which the slides were placed in 10 mM sodium citrate buffer (pH 6), boiled at 700 W in a microwave, and left at 120 W for another 9 min. After cooling down, incubation in increasing ethanol series and a short period of air-drying, the hybridization with the PNA FISH probes was performed. For this, tissue sections were incubated with 0.1 µM of a Cy3-labeled telomere probe (CCCTAA)_3_ (TelC-Cy3, Panagene). In experiments where the centromeres were also visualized, 0.1 µM of a FAM-labeled CenpB PNA probe (ATTCGTTGGAAACGGGA) was added at the same time. The hybridization took place in 70% formamide, 10 mM Tris-HCl, pH 7.5, 0.1 µg/ml salmon sperm. First, slides were denatured at 84°C for 5 min and then left overnight at room temperature in a wet chamber for hybridization. Next, slides were washed three times for 15 min in PNA wash buffer, followed by three 5 min-washes in PBST, and incubation with an anti-PML antibody (1:100, PG-M3, sc-966) in PBS overnight at 4°C in a wet chamber. Finally, the slides were washed with PBST, incubated with the secondary antibody (here: anti-mouse IgG coupled to Alexa647) for 1 h at RT, again washed with PBST and embedded with Prolong including DAPI.

### 2.4. Image acquisition

Wide-field images were acquired with a fully automated Olympus IX81 ScanR screening microscope. A magnification of 10x (Olympus UPlanSApo, NA 0.4) was chosen, resulting in a field of view of 866 µm x 660 µm. The sample was illuminated with a 150 W Hg/Xe mixed gas arc burner in combination with appropriate filter combinations for DAPI and Cy3, respectively. For the TMAs, 725 (29 x 25) adjacent fields of view were recorded for cell selection, with overlapping areas of 30 µm in x- and y-direction in order to enable automatic image stitching. In total an area of ~3.8 cm^2^ was imaged. For confocal imaging a Leica TCS SP5 point scanning confocal microscope with additional MatrixScanner software was used. In all cases, a 63x objective lens (Leica HCX PL APO, NA 1.40) was employed. Scan speed, pixel size, field of view, z-stacks and channels were set for each sample appropriately; typical values for high-resolved imaging of previously identified regions of interest (ROIs) were 200 Hz scan speed, 96 nm pixel size at a field of view of 256 x 256 pixels, 41 axial layers at 250 nm spacing and 3 color channels with 2x frame averaging. Single molecule localization microscopy (SMLM) was performed with the same Leica TCS SP5 microscope used for confocal imaging, equipped with an additional wide-field unit. For illumination, a 647 nm diode laser with 140 mW maximum power (Luxx, Omicron, Rodgau-Dudenhofen, Germany) and a 561 nm solid state laser with 156 mW maximum power (Jive, Cobolt, Solna, Sweden) were used. The laser beam path was additionally coupled into the microscope through the back illumination port by a switchable mirror. Highly inclined and laminated optical sheet (HILO) illumination of the sample was controlled by an adjustable mirror. Images were recorded by an sCMOS camera (Hamamatsu Orca Flash 4.0) at 10 ms integration time and 5 000 frames per dataset with each maximum laser power in order to induce photoswitching. From the total field of view of 2048 x 2048, a sub-region of 200 x 200 pixel (20 x 20 µm^2^) was acquired. Prior to SMLM imaging, a wide-field image was acquired at low laser intensities. The area illuminated by high laser power was restricted to the target cell by an adaptable aperture in the illumination beam path in order to protect surrounding structures from photobleaching. SMLM datasets were reconstructed using the ImageJ plugin ThunderSTORM [33].

### 2.5. 3D image analysis

To adapt our previous work on the automatic extraction of telomere features from cell line samples to tissue sections, an advanced 3D model-based image analysis approach was developed. It accounts for the elevated levels of tissue sample heterogeneity, image artifacts and significant background levels (Fig. 4A) [31, 34]. The approach supports multiple microscopy channels, where depending on the experiment a maximum of four 3D channels was used (i.e., centromeres, telomeres, PML-NBs and DAPI staining). To overcome artifacts caused by false positive segmentation such as autofluorescent erythrocytes (Fig. 4E), first a 3D artifact detection and segmentation step is performed. Artifacts were determined in the telomere channel and segmented in 3D by performing noise reduction (3D Gaussian filter) followed by automatic thresholding (intermodes scheme), different morphological operations, and object filters. The inverted binary segmentation result was used as a mask for subsequent segmentation of the cell nuclei in the DAPI channel. For the 3D segmentation of the cell nuclei an approach similar to ref. [31] was used. It is based on different automatic thresholding schemes (e.g., Otsu, entropy-based) where the artifact mask was included. The nuclei segmentation result defines a mask for the analysis of all other channels, and it serves for normalization of analysis results with respect to the volume of the cell nuclei. Depending on the experiment, a further labeling step was carried out to enable a cell-based analysis.

For automatic quantification of relevant spots in the telomere, PML and centromere channels, a series of 3D spot detection, 3D model-based segmentation, and spot filtering was performed. To automatically detect spot candidates, different image analysis operations were applied to the 3D image data to obtain (coarse) center positions of the spots: For noise reduction we used either a 3D Gaussian filter or a 3D Laplacian of Gaussian filter. For suppressing the image background all intensity values below a certain threshold value, which was automatically computed for each image from the image histogram based on the region of the cell nuclei [34], were clipped. Finally, we performed a local maxima search within cubic 3D ROIs, where the local maxima represent the spot candidates. In the second step, each detected spot candidate was quantified based on 3D least-squares model fitting using a 3D Gaussian parametric intensity model. A 3D Gaussian model well represents the 3D intensity profile of the considered spots. The model reads

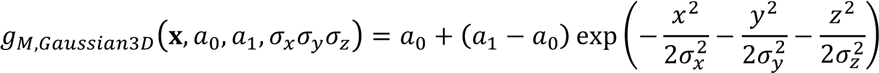

where **x** is a 3D position, *a*_0_ and *a*_1_ denote the local background and peak intensity levels, respectively, and *σ_x_, σ_y_*, and *σ_z_* are the standard deviations that represent the three semiaxes describing the ellipsoidal shape of the spots. For further details see ref. [34]. By fitting parametric intensity models a subvoxel resolution of the model parameters including the 3D position and semi-axes is obtained. To enable accurate and robust fitting of the variably sized telomere spots, the size of the 3D ROI used for model fitting was automatically determined for each spot by a multiscale scheme similar to that described in ref. [35] (Fig. 4B-D). Here the spot size is coarsely determined by analyzing the results of a 3D Hessian-based blob detector at different image scales. In contrast to the scheme in ref. [35], which has been designed for larger spots of irregular shape, we here account for the heterogeneous spot sizes by adapting the detector response. Finally, based on the quantified model parameters, different spot filters were applied to the fitting results to exclude invalid spots, e.g. those that significantly deviate from a typical spot regarding their shape or size or that displayed a low image contrast as they likely represent noise. Since the image statistics vary among different images, a threshold for the contrast based on the image histogram in the region of a spot was computed. For calculating the integrated intensity of each fitted spot, an analytic formula was derived that integrates the intensities of the 3D Gaussian model over the ellipsoidal volume. The analytic solution of the triple integral reads

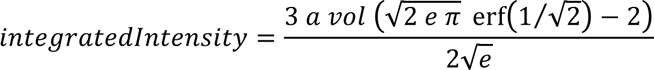

where *a* = (*a*_1_ – *a*_2_) denotes the spot contrast, *vol* = *σ_x_σ_y_σ_z_* 4π/3 is the ellipsoidal volume of spot, and erf denotes the error function. The integrated intensity is computed from the fitted model parameters, i.e., the noisy image intensity values were *not* directly used.

To determine cell-based co-occurrences as well as 3D colocalizations the results of pairs of spot channels were combined. The accessibility of the PNA FISH probes (see section 3.2, Fig. 2B) can be assessed with a 3D cell-based co-occurrence analysis that quantifies the number of centromeres and telomeres per cell nucleus. For the quantification of colocalizations between telomeres and PML-NBs the 3D spot geometries were obtained with subvoxel accuracy from model fitting. These data were subsequently used to determine whether two spots from different channels spatially overlap in 3D [31] (Fig. 4F, G). Finally, results of all channels and analysis steps are combined and all results for each single sample were summarized and further statistics computed. The telomere length distribution was quantified for each sample based on the histogram of the integrated intensities of all telomere spots, and normalized with respect to the volume of DAPI staining.

For each experiment the same set of parameters was applied and all images were analyzed in batch mode (running twelve jobs in parallel). The computation time for a single multichannel 3D image was about 30-40 s, and a full TMA required about 1 hour on an Intel Xeon CPU with 2.67 GHz running Linux. For the exemplary application shown here a total 2583 multichannel 3D images from prostate cancer TMAs and 247 such images from several pedGBM sections were evaluated.

## 3. Results and Discussion

### 3.1. An integrated workflow for the automated fluorescence microscopy and analysis of telomere features in primary FFPE tumor tissue sections

The overall approach for the 3D imaging of FFPE tumor tissues described here is illustrated in Fig. 1. It consists of three technically challenging parts: The first part involves the multi-color staining of the samples with three types of staining that are informative for the TMM analysis. (i) FISH of telomeres to identify their number and length distribution. In addition, FISH staining of centromeres provides a valuable control. It indicates whether the absence of a telomere FISH signal indeed reflects very short telomeres or simply the failure to hybridize the probe in a certain part of an image. (ii) Immunofluorescence (IF) of PML protein, which can be evaluated in terms of its colocalization with telomeres as a marker for ALT. (iii) Counterstaining of the DNA with DAPI to identify nuclei.

**Fig. 1.**
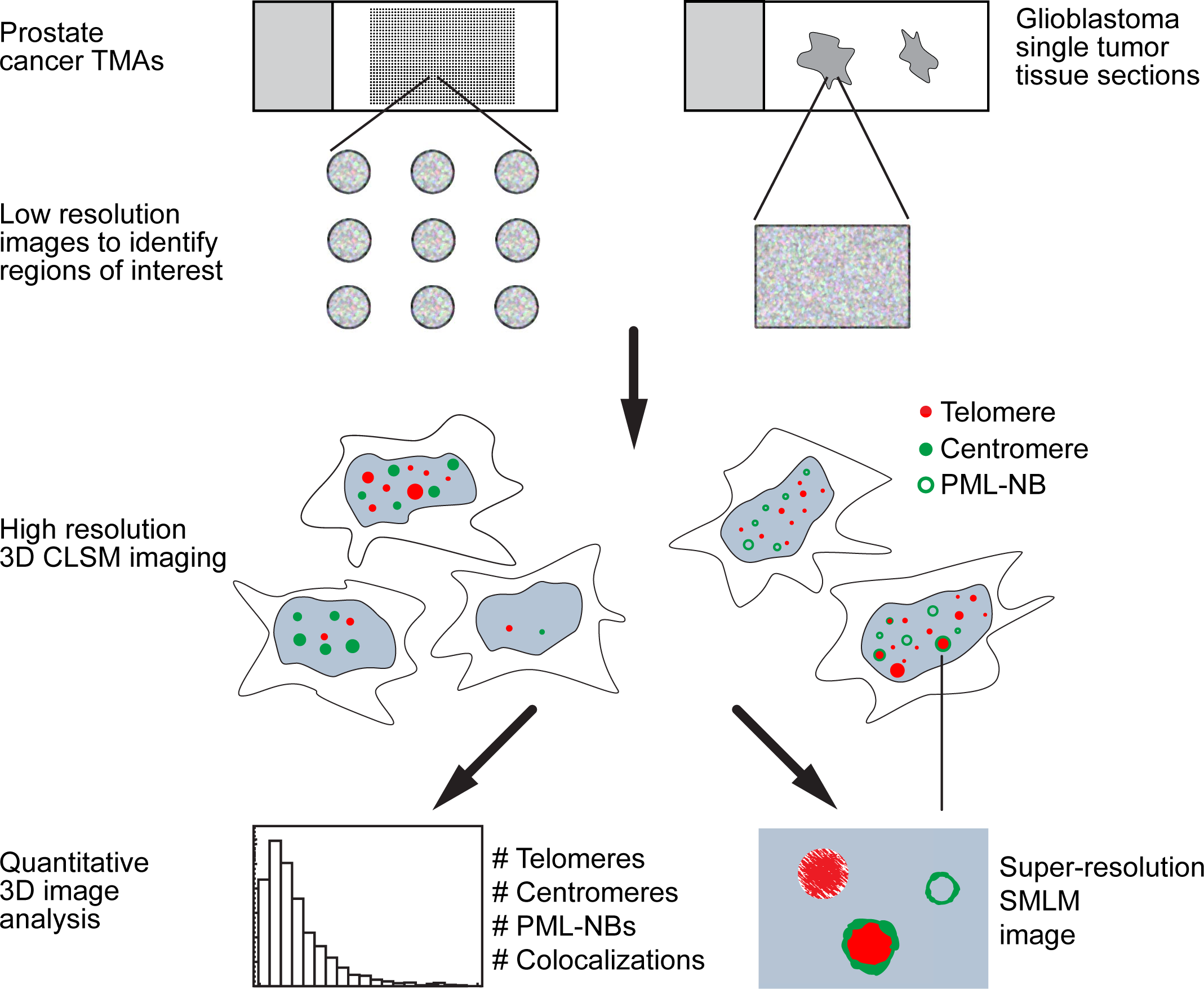
Schematic overview of workflow. The 3D imaging and analysis of FFPE tumor tissues either on TMAs or as single sample sections consists of the three main parts: sample staining, image acquisition and image analysis. These steps are illustrated in detail in Figs. 2-4. Briefly, the fluorescence labeling comprises FISH of telomeric and centromeric repeat sequences, immunofluorescence of PML protein and counterstaining of the DNA with DAPI. The image acquisition involves a combination of 2D imaging with a wide-field setup followed by targeted CLSM imaging with an optional zoom-in step by super-resolution microscopy (SMLM). In the 3D image analysis, subnuclear structures are identified from 3D spot detection to quantify number, size, signal intensity and co-occurence of telomeres, centromeres, and PML-NBs. In this manner, crucial information on the active TMM in a given tumor sample is obtained.

**Fig. 2.**
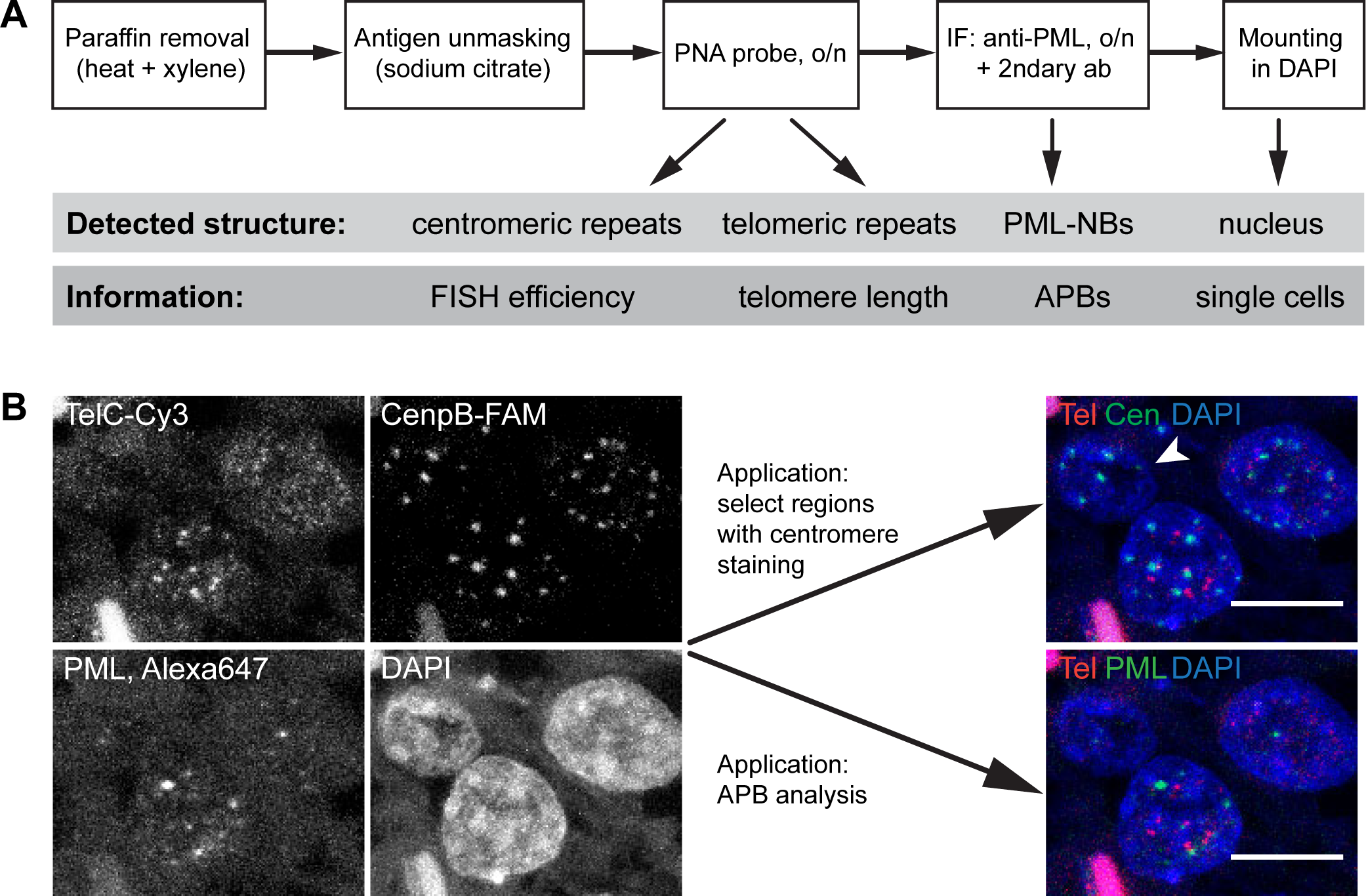
Staining of nuclear structures in FFPE sections. **A.** Schematic workflow for the staining procedure of FFPE tissue sections to detect telomeric and centromeric repeats by PNA FISH probes as well as PML-NBs by immunofluorescence against the PML protein. Additionally, the nucleus is visualized by DAPI staining. The detection of these nuclear structures allows the discrimination of single cells and provides information about the efficiency of the FISH probe hybridization, the length of the telomeres, and the presence of APBs. **B.** Representative CLSM image of a pedGBM tissue section that has been treated with PNA probes against the telomeric repeat sequence (labeled with Cy3) and against centromeric repeats (labeled with FAM) and additionally stained for PML protein (labeled with Alexa647). The nuclei are detected by DAPI. Two merged images are shown. The upper one shows a merge of the telomere (red), the centromere (green), and the DAPI (blue) signal. The arrow indicates a nucleus that was accessible for the FISH probes as is obvious from the detection of the centromeres but that did not display any signal from the telomere FISH probe. Accordingly, this cell can be classified as one that has very short telomeres that are not detectable using telomere FISH as opposed to not being accessible for the FISH probe. Below is a merged image of the telomere channel (red), the PML IF (green) and DAPI (blue). Using this combination the presence of APBs can be analyzed. All images are maximum projections. Scale bars, 10 µm.

The second part of the workflow comprises the image acquisition: (i) First, the whole tissue sample is imaged in 2D with a wide-field setup. (ii) Images are analyzed on-the-fly to identify regions of interest, e.g. nuclei showing DAPI and/or telomere signal. (iii) Information on the corresponding positions is fed back to the microscope. (iv) High-resolution multicolor confocal fluorescence 3D images are acquired for an in-depth automated analysis of subcellular structures at the regions of interest. (v) An optional zoom-in step by photoactivated localization microscopy can be applied for analyzing the structure of selected parts of the image at a resolution that is higher by an order of magnitude.

The third part of the workflow consists of an automated 3D image analysis: (i) A 3D artifact detection and segmentation step removes artifacts like autofluorescent signal from contaminating cells or debris. (ii) Regions of interest are selected via a 3D segmentation of the cell nuclei. (iii) Subnuclear structures are identified from 3D spot detection and quantification. (iv) A 3D cell-based co-occurrence analysis can be performed by analyzing the number of quantified centromeres and telomeres within each cell nucleus. (v) Via a 3D colocalization analysis, APBs are identified from co-occurring telomere and PML signals.

### 3.2. Combined immunofluorescence and FISH staining for detection of ALT positive tumors using confocal laser scanning microscopy

Usually the ALT status of a fixed tumor tissue is determined by telomere FISH alone. However, a few issues should be considered. Different fixation conditions used for different tumor samples as well as heterogeneous results within one tissue sample can make some areas on a slide more accessible for the PNA FISH probe than others. This is particularly important when comparing several tumor samples with respect to their telomere lengths. Usually, it is assumed that a single sample or all the tissue cores on a given TMA slide have minimal technical variations since they are stained in the same manner. Thus, the lack of a telomeric signal would indicate the presence of only very short and thus undetectable telomeres. Since the absence of a telomere FISH signal could alternatively be due to the inaccessibility of this particular cell for the FISH probe. To control for this issue, a second PNA FISH probe can be applied [25]. For comparison this should also be directed against a repetitive sequence in the genome, such as centromeric repeats. The detection of a centromere FISH signal in a given cell indicates that technically the FISH probe can access this nucleus and a lack of telomere FISH signal would indeed be due to the presence of very short telomeres. If no centromere signal is observed, this would argue that none of the probes could access the nucleus. Accordingly, such a cell should be omitted from the analysis. Secondly, it is problematic to rely only on the telomere signal to identify a tissue as ALT positive. It is widely accepted that tumor cells and cell lines that use the ALT mechanism have a very heterogeneous telomere length distribution within a cell population and also within a single cell. However, in most studies telomere FISH on tissues the telomere length distribution is not determined but only the presence of very strong ultra-bright foci is evaluated. These foci are used as an ALT marker despite the uncertainty about their biological source and function. Thus, for a more reliable characterization of a tissue section with respect to its TMM, the presence of APBs should be assessed, since these nuclear subcompartments are specific for ALT positive cells [9]. By simultaneous detection of telomeres and PML-NBs as well as an automatic quantification of the colocalizations the number of APBs can be determined [30, 31].

To address these issues we stained FFPE tissue sections of pedGBM samples with FISH PNA probes against telomeric and centromeric repeats as well as with IF against the PML protein. Images were acquired by CLSM and analyzed automatically with respect to the telomere length distribution and frequency of APBs. The workflow of the sample preparation is illustrated in Fig. 2A. Essentially a protocol similar to that given in ref. [25] was used as described in the Material and Methods section. As indicated in Fig. 2A, an appropriate combination of dyes allows the simultaneous detection of centromeres, telomeres, and PML-NBs in single cells. In Fig. 2B some representative CLSM images are shown. The combination of centromere and telomere signal is analyzed for quantifying the telomere length distribution in a given sample (Fig. 2B, top merge image). This combination of readouts is useful to assess the efficiency of the FISH probes and to evaluate whether the lack of a telomere signal is indeed due to the presence of very short telomeres or attributed to technical issues. Additionally, the telomere signal can be analyzed in relation to a PML-NB staining (Fig. 2B, bottom merge image). With this combination of stainings the frequency of APBs can be evaluated. From the number of APB the TMM of tumor cells can be inferred as described below in further detail.

### 3.3. 3D Targeted Imaging

In order to identify and relocate ROIs on a TMA, two routines have been implemented. In one approach, ROIs are automatically preselected within the images by a wide-field-screen (Fig. 3A-C). The other routine uses on-the-fly switching of the optical configuration upon automatic identification of an ROI (Fig. 3D, E). In the first case, a complete TMA was scanned on an Olympus IX81 microscope with a magnification of 10x. Images were recorded in the DAPI color channel with an overlap of 30 µm in x and y direction to ensure complete coverage and to enable stitching of adjacent images (Fig. 3C, left). The resulting images were processed within the ROIs using KNIME (Konstanz Information Miner, www.knime.org) and the KNIME image processing plugins (KNIP). A background subtraction, a combination of global and local thresholding and subsequent connected component analysis was applied. The results were filtered for size and intensity in order to extract structures that resemble cells. The global position of each ROI on the TMA was retrieved from the stage position of each image (image metadata) together with the position of the ROI within the respective image. Positions were assigned to the associated spots of the TMA based on the regular pattern of these spots. A distance filter was applied to remove ROIs closer than 25 µm in order to avoid overlapping imaging. The corresponding KNIME workflow is depicted in Fig. 3B (upper part). The sample was then transferred to a Leica TCS SP5 confocal microscope and one image of the DAPI staining was acquired in the top left, top right and lower right spot. These three images were used to match and transfer the coordinates of the ROIs from the wide-field to the confocal microscope (Fig. 3B, lower part). A list of the ROIs within the coordinate system of the stage of the confocal microscope was generated, which was automatically processed by the microscope. At each ROI position, a confocal stack with three colors (Cy3-Telomere, FAM-Centromere, DAPI-nucleus) and 41 z-planes (distance 0.25 µm) was acquired (Fig. 3C, most right).

**Fig. 3.**
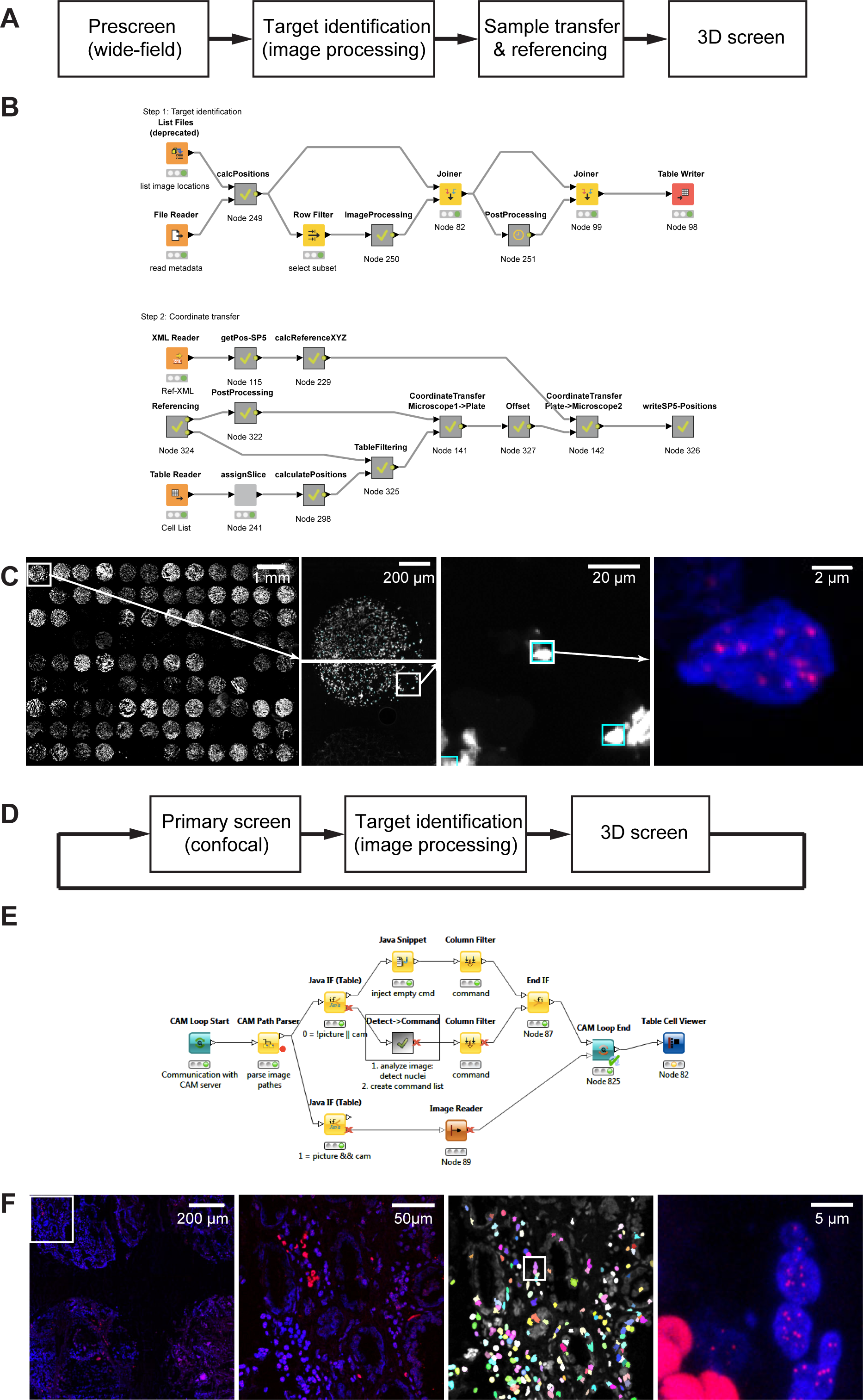
3D Targeted Imaging (3D-TIM) **A.** Schematic workflow for sequential imaging on different scales. First, regions of interest are identified in a pre-screen. Then, the sample is transferred to a second microscope and after a referencing step the previously identified regions can be addressed and acquired in 3D with higher resolution and more color channels. **B.**KNIME implementation of the workflow in A. **C.** Example images. On the left, the single tissue spots of a TMA can be seen. Next to it, the two wide-field images acquired of the first spot are shown. These were processed in order to identify the ROIs as described in the text. On the right side, the same region is shown as imaged by CLSM after sample transfer and referencing (maximum projection of 41 axial layers). **D.** Schematic workflow for integrated imaging on different scales. In each image of a primary screen, regions of interest are identified. These positions are directly fed back to the microscope and a high-resolution z-stack is acquired for each region. Afterwards, the next scan field of the primary screen is acquired. **E.** KNIME implementation of the workflow in D. **F.** Example images. On the left, the single spots of a TMA can be seen. Next to it, a two-color image of the highlighted region of the confocal prescreen is shown. This image is directly processed in order to find ROIs (i.e. cellular structures comprising five or more telomere spots) indicated in random colors. On the right side, one region is shown as imaged by the subsequent 3D screen (maximum projection of 41 axial layers).

**Fig. 4.**
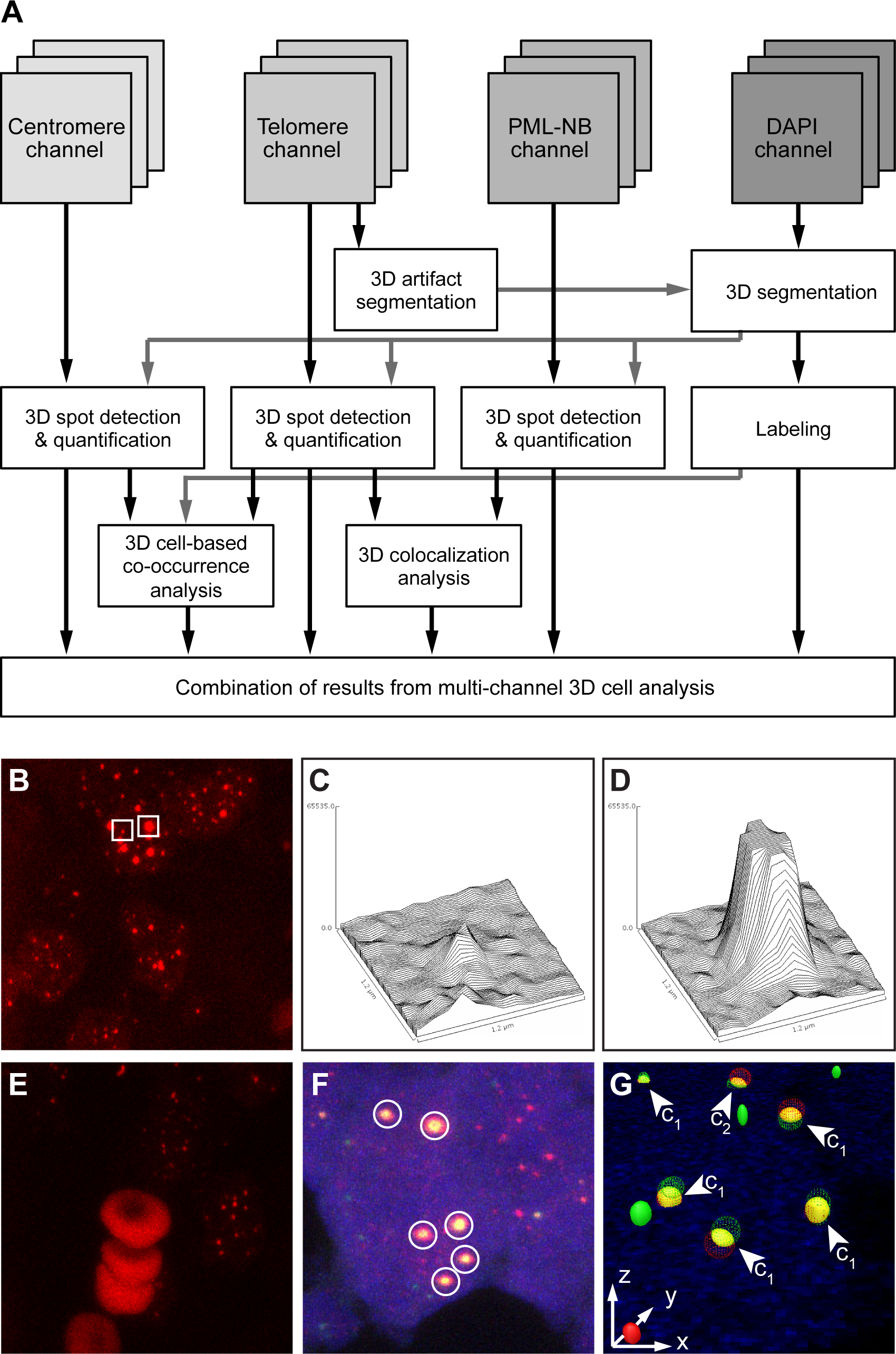
Automated 3D image analysis of telomere features from tissues. **A.** Schematic workflow for 3D quantitative image analysis of 3D multichannel microscopy images. Subnuclear structures are identified using 3D spot detection after 3D segmentation of DAPI stained regions in the different color channels. Subsequently the number, size and signal intensity of telomeres, centromeres and PML-NBs are quantified using a 3D model-based approach. Finally, 3D cell-based co-occurrence and co-localization analyses are conducted. **B.** Example image of the telomere channel of a prostate cancer TMA showing telomere spots of heterogeneous sizes. **C, D.** 2D intensity plots of a small telomere spot (C) and an ultra-bright telomere spot (D) (both highlighted in B). **E**. Example image of the telomere channel from the TMA image acquisition depicting large artifacts (erythrocytes). **F.** Overlay of telomere, PML, and DAPI channels of a pedGBM sample, where six colocalizations of PML bodies with large telomeres are highlighted. **G.** 3D visualization of the quantified geometry of the telomeres, PML bodies, and colocalizations for the image in F with marked colocalization classes c1 and c2. Images in B, C, and D are maximum intensity projections.

For the second image acquisition routine, a TMA was scanned in 2D at medium resolution on a confocal Leica SP5 microscope using the Leica MatrixScreener software (Fig. 3F, left). This software provides a server-client interface (termed ‘computer aided microscopy’, CAM), by which the scan routine can be influenced and altered. The scan was embedded into a KNIME workflow loop (Fig. 3E). Special loop start and loop end nodes for addressing the CAM-interface have been developed in collaboration with KNIME and Leica. In each step of the loop, one scan field was imaged and directly processed within KNIME via the KNIME image processing plugins (KNIP). Cells were identified based on the DAPI staining, and single telomeric spots within these cells were counted based on the Cy3 signal. Nuclear structures containing five or more telomeric spots were considered as candidates for high-resolution 3D imaging. The positions of these candidates were collected and fed back to the microscope via the KNIME-CAM-interface. Upon receiving these positions, a second high-resolution 3D routine was started immediately at the microscope that acquired those candidates. Afterwards the next iteration of the loop was started and the next scan field was recorded.

The first routine can be used if a set of images of a specific sample already exists and specific subregions are targeted for higher resolution image acquisition. These subregions can be identified manually by investigating the whole dataset and marking structures of interest. Alternatively, automatic segmentation and classification routines can be applied to the whole dataset, subpopulations can be identified and representatives of these populations can be acquired with higher resolution in 3D or in additional color channels. If the phenotype of the structures to be acquired is known beforehand, the second routine can be used. The microscope then acts as a trigger system, which can switch the acquisition mode upon registration of a predefined event (detection of a structure of interest, rare cell cycle event, etc.), acquires images of the desired resolution, and afterwards continues the screen.

### 3.4. Extraction of telomere features from fixed tumor sections via automated 3D image analysis

To extract telomere features from fixed tumor sections, a robust 3D model-based image analysis approach described in detail in the Material and Methods section was developed (Fig. 4). This approach includes an artifact detection step to exclude regions of artifacts from further analysis (e.g. fragments of red blood cells), a cell segmentation step, and 3D model-based spot quantification. The approach is fully automated without the need of manual parameter adjustments for individual images. For the application shown here, three fixed sets of parameters were used: one for the analysis of tissue spots from TMAs of prostate cancer and two for the analysis of pedGBM tissues. Common parameter settings for all images included the chosen parametric model for spot quantification (anisotropic 3D Gaussian intensity model), the initial model parameters, the range of 3D ROI sizes used for model fitting (radii of 3-10 voxels), and the 3D ROI size for local maxima search (5 × 5 × 5 voxels). In contrast, the parameter settings for 3D spot detection, spot filters, and 3D DAPI segmentation were adapted to the intensity distribution obtained for the staining of a given TMA or tissue sample. For the spot detection in the TMA images, a 3D Gaussian filter with isotropic smoothing (σ=1 voxel) was used for noise reduction followed by intensity clipping using relative factors of *c*=2 for telomeres and *c*=4 for centromeres (see [34] for details). In the images from the pedGBM sections, a 3D Laplacian of Gaussian filter with anisotropic smoothing (σ_x,y_=1.5 and σ_z_=1 voxels) was used as well as relative factors of *c*=14 or *c*=16 for telomeres, centromeres, and PML-NBs. Furthermore, spots were filtered using the minimum spot contrast with relative factors of *s*=2 (prostate cancer images) and *s*=3 or *s*=4 (glioblastoma images) with respect to the standard deviation of the image histogram. In order to segment the DAPI signals Otsu thresholding was employed with one threshold (TMA images) as well as Otsu thresholding with two thresholds or maximum entropy thresholding (pedGBM images).

As an example, Fig. 4G shows a 3D visualization of the 3D segmentation results of telomeres (red) and PML-NBs (green) of one multichannel microscopy image of the glioblastoma sample shown in Fig. 4F. In our approach a 3D geometric description of the telomere and PML-NBs spots is obtained. From this parameterization, colocalizations between these spots are detected and classified automatically (Fig. 4G). Two different classes of colocalizations were defined: In class 1 (c1) the center of one spot lies within the other spot. The class 2 (c2) colocalization is a less stringent definition that only requires an overlap of the spots. Note that with standard approaches, colocalizations are usually determined based on color overlays of the red and green channels (see discussion in ref. [34]). Such color overlays strongly depend on the image contrast and background signal in the different channels. They were not applicable for the tissue samples due to the high heterogeneity of intensity variations and image artifacts. In contrast, by using a 3D geometry-based approach colocalizations could be determined robustly in the presence of image contrast variations.

### 3.5. 3D-TIM of telomeres on a prostate cancer TMA reveals differences in the telomere length distribution between tissue cores

In order to illustrate the automated image acquisition and analysis we evaluated 45 prostate cancer tumor sections on TMAs with respect to their telomere signals detected with a Cy3-labeled FISH probe. The telomere length distributions of four representative tissue cores from the TMA are presented in Fig. 5. The amount of hybridized telomere probe is directly proportional to the number of telomeric repeats and hence the signal intensity is a direct indication of the telomere length. The distribution of the telomere intensities from different prostate cancer biopsies revealed the heterogeneity of the telomere length distribution between tissue spots. For example, the maxima of the distribution for II and III were shifted towards longer telomeres as compared to I and IV, which had the highest frequency at the lowest intensity bin (Fig. 5). In addition, samples III and IV had a higher fraction of very long telomeres than I and II. This type of quantitative information cannot be obtained from the commonly performed manual examination of wide-field images. The comparison of the frequencies of telomere intensities on a cell-to-cell basis between different tissues requires an automated segmentation of individual DAPI stained nuclei in FFPE tissues. This step is technically challenging as nuclei in very close vicinity often cannot be well separated from each other due to low image contrast and significant background signal. To overcome this problem, normalization to a measured DAPI volume has been implemented (Fig. 5B). Currently, one caveat here are relatively high background DAPI signals in some of the tissue sections. One example is depicted in histogram IV where an abnormally high DAPI volume was measured. In order to address this issue the ratio of the centromere FISH to DAPI signal can be determined (#centro/1000 µm^3^ in Fig. 5). A low value of this parameter is indicative of technical problems in the region evaluated: Either the measured DAPI signal has an unusually high background or the FISH probe hybridization was not efficient on this particular tissue spot. It is advantageous to conduct this analysis with the centromere probe. The number of centromeres per nuclei should reflect the chromosome content of the cell, whereas the detectable telomere signals is affected by the efficiency to telomere maintenance and cell proliferation. Thus, a low centromere/DAPI ratio represents an informative quality parameter to identify tissue regions that should be excluded from the quantitative analysis of the telomere length distribution due to an inefficient FISH reaction. In contrast regions with normal centromere/DAPI ration but the low telomere signal can be assigned to cells with very short telomeres. In the future, this analysis can be further improved by reducing the unspecific DAPI signal that is prominent in some of the spots, for example by using different DNA staining reagents. This will also help to implement a proper automatic segmentation of the nuclei and allow for a cell-based analysis of the FISH signals. Further improvements will include the use of a gradient-flow tracking approach for the cell-based nucleus segmentation as well as an extended artifact detection step that exploits all microscopy channels.

**Fig. 5.**
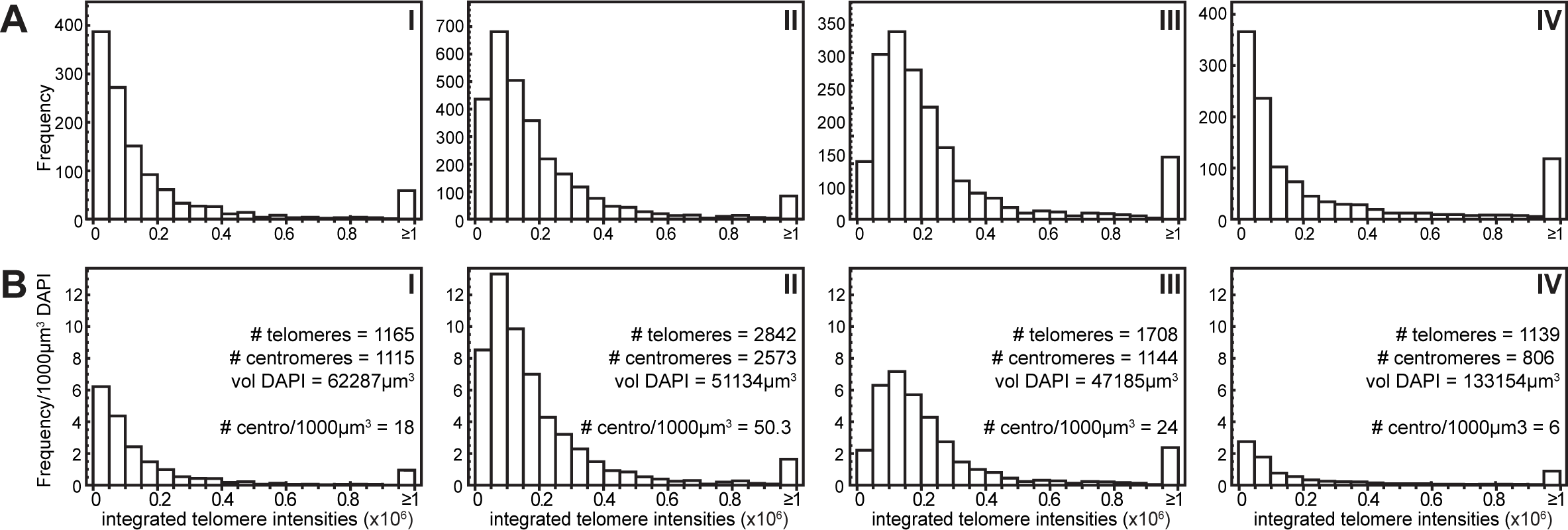
Distribution of telomere signal intensities in different samples of a prostate cancer TMA. Exemplary results from analyses of four samples (I, II, III, IV) from a prostate cancer TMA that was stained using telomeric and centromeric FISH probes. Images were acquired by the 3D-TIM mode, centromere and telomere signals were automatically counted, and telomere signals were analyzed with respect to their integrated intensity. Histograms were generated using 20 bins covering intensity units from 0 to 1x10^6^ (A.U.). Values >1x10^6^ are summarized in the last bin. **A.** Distribution of the absolute frequencies of integrated telomere intensities in four different tumor tissue cores from the TMA, **B.** Same analysis as in A, only frequencies were normalized to the detected DAPI signal.

### 3.6. Quantitative confocal microscopy analysis of tissue sections can be used to determine the active TMM

A functional TMM is crucial for unlimited proliferation, a hallmark of tumor cells. Accordingly, several recent studies concluded that the type of active TMM can be exploited for patient stratification and prognosis [3, 4, 19–22]. In order to determine the active TMM in primary tumor samples, several characteristic features of ALT can be addressed by different methods depending on the material available. One very reliable marker of ALT activity is the presence of ECTRs that can be visualized by the C-circle assay [15]. This assay, however, requires DNA of high quality isolated from frozen tumor tissues. For FFPE-fixed tissues, the detection of APBs by simultaneous PML-IF and FISH hybridization of telomeres is well suited to determine the ALT status (Fig. 2, Fig. 6). FFPE sections from three different pediatric glioblastoma samples (labeled with I, II, and III) were stained and imaged (Fig. 6A, 6D). Then an automated 3D image analysis was applied to quantify their telomere length distribution (Fig. 6B, C) as well as the colocalization of telomeres and PML-NBs (Fig. 6C). Of the three tested pedGBM samples, one tumor was identified as ALT positive (Fig. 6, III) as this tissue section revealed a higher frequency of colocalizations between PML protein and telomeres (11.0% colocalizations for all telomeres in III as opposed to 4.9% in I and 4.4% in II). Notably, this APB-positive tumor sample also shows a distinct telomere length intensity profile (Fig. 6B, D) with a high portion of more intense telomere signals compared to the APB-negative tumors. Indeed, 13% of the detected telomere signals exhibited an integrated intensity of ≥ 1,000,000 (A.U.) compared with only 1% or 4% of telomeres with such strong signals in the other two tested tumor samples. ALT positive cells frequently show an elevated telomere repeat content. Most likely this is reflected by the high number of strong signals obtained from the telomere FISH probe in this case (Fig. 6B-D). This type of analysis can be extended by the cell-based analysis of a concomitantly applied centromere-specific FISH probe in order to get a more precise understanding of the telomere length distribution of the different samples as explained above.

**Fig. 6.**
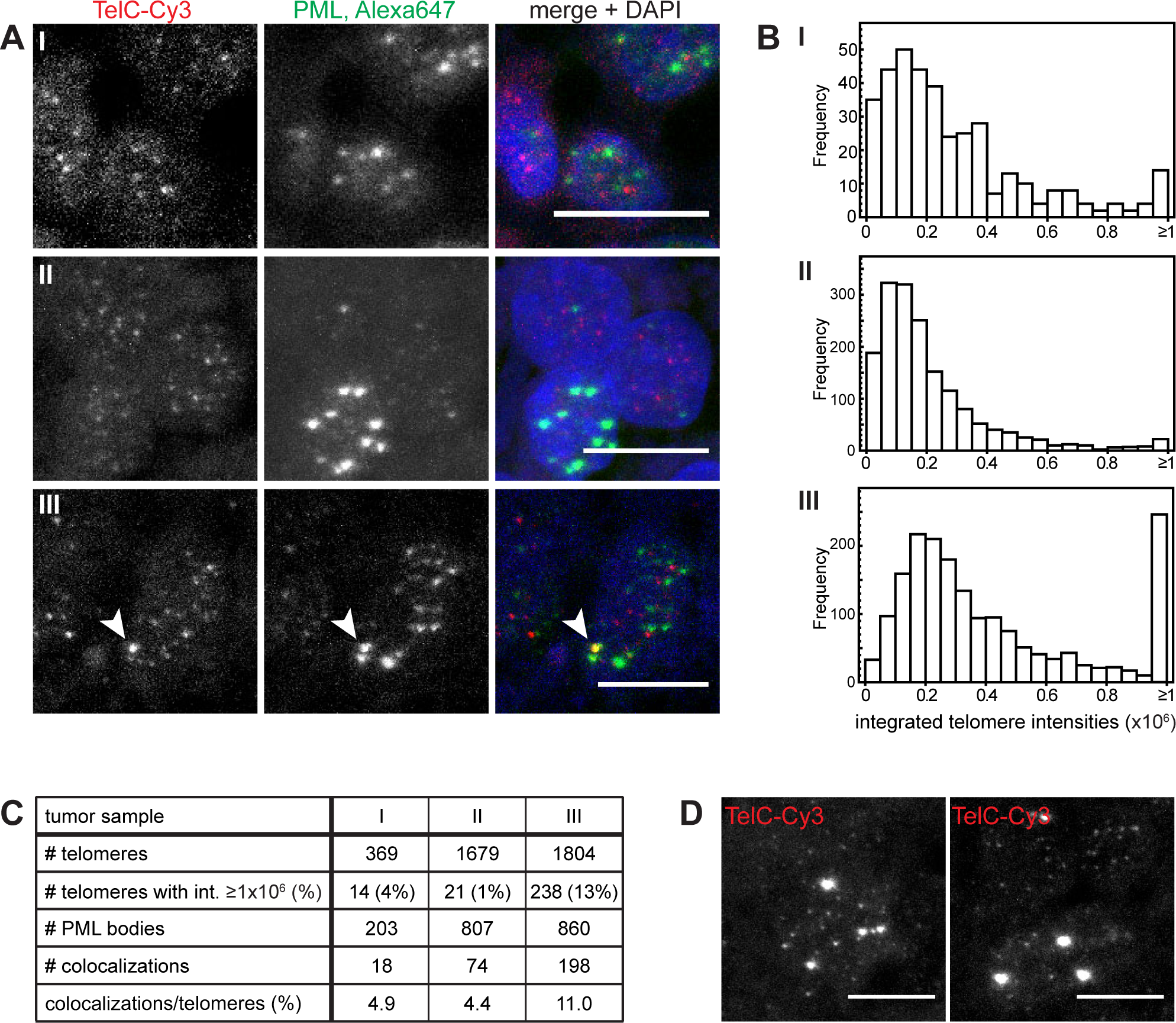
Identification of the ALT status of pedGBM samples from FFPE tissues. **A.** Representative CLSM images from tissue sections of three pediatric glioblastoma samples (I, II, III) stained for telomeres and PML protein. A colocalizing telomere and PML signal in tissue section III is indicated by an arrow. **B.** The distributions of telomere lengths in the analyzed tumors are visualized as histograms of the integrated telomere intensities as determined from the respective tissue sections by automated 3D image analysis of the CLSM images. Note the high number of telomeres with an integrated intensity of ≥ 1x 10^6^ (A.U.) in the APB-containing tumor III compared to the other tumor samples that do not show ALT-specific features (see also Fig. 4D). **C.** Overview of results from automated image analysis. Telomeres, PML signals, and colocalizations were detected and quantified automatically in the 3D images from the CLSM acquisition as shown in A. Tumor sample III exhibits an increased number of telomeres that colocalize with a PML-NB indicating an active ALT mechanism in this tumor. **D.** Examples of cells from tumor section III with very bright telomere spots as detected by the telomere-specific PNA FISH probe. All images are maximum projections. Scale bars, 10 µm.

### 3.7. An integrated super-resolution mode reveals more details on telomeres and PML-NBs in tumor tissue sections

In further analyses, PML-NBs and telomeres in FFPE-fixed glioblastoma tissue sections were examined using single molecule localization microscopy (SMLM, Fig. 7 and Material and Methods). In super-resolution techniques based on photoswitching of single fluorescent molecules, only a changing small random subset of the fluorophores emits light in a given image frame. Thus, the signals from individual fluorophores can be isolated and fitted by a model function to determine their location with an accuracy in the range of a few nanometers. Usually several thousand image frames are acquired, so that the labeled structures can be reconstructed afterwards based on the fit coordinates [36–40]. Using such an approach, PML-NBs labeled with Alexa647 by immunofluorescence against the PML protein could be resolved in primary tissues and additional information compared to wide-field and CLSM images was inferred (Fig. 7A-C). First, the sizes of the subnuclear structures can be determined in a more precise manner. Here, the measured sizes of the PML-NBs in the super-resolution mode varied from about 100 nm to 800 nm. Occasionally, hollow structures with an estimated thickness of the outer shell of about 100 nm were observed (Fig. 7C, I, III). This is in good agreement with previously described characteristics for PML-NBs in U2OS and HeLa cell lines detected with 4Pi super-resolution microscopy [41]. In accordance with previous reports on the structure of these nuclear bodies in cell lines, we could detect an irregular distribution of PML protein on the outer shell in the primary tumor tissue sections [41]. Besides the larger structures, several smaller spots of enriched PML signal were also detected (Fig. 7C, II and IV). They might represent stress-induced PML microbodies or nuclear microspeckled structures as have been described before [42], but this needs further confirmation. Notably, the Cy3 dye, which previously has been reported to undergo photoswitching but produce only low quality reconstruction images [43], could also be used for the localization approach in the tissue sections (Fig. 7B, D). Using the super-resolution mode on the Cy3-labeled telomeres we were able to detect an increased number of telomere signals (Fig. 7A, B). Further, the sizes of telomere signals can be assessed more precisely using super-resolving methods for short telomeres (Fig. 7D, F) as with confocal microscopy when the extension of the telomeric signal is at the border or even below the resolution limit. In the future this could be exploited for a more accurate determination of the telomere length distribution. With simultaneous use of Alexa647-labeled PML protein and the Cy3-labeled telomere FISH probe, coordinate based analyses of the single molecule distributions become feasible [44–46]. Thus, this approach can also be applied to get further insight into the structure of APBs in ALT positive tumor tissues and to compare these findings to previous studies on APBs in cell culture models.

**Fig. 7.**
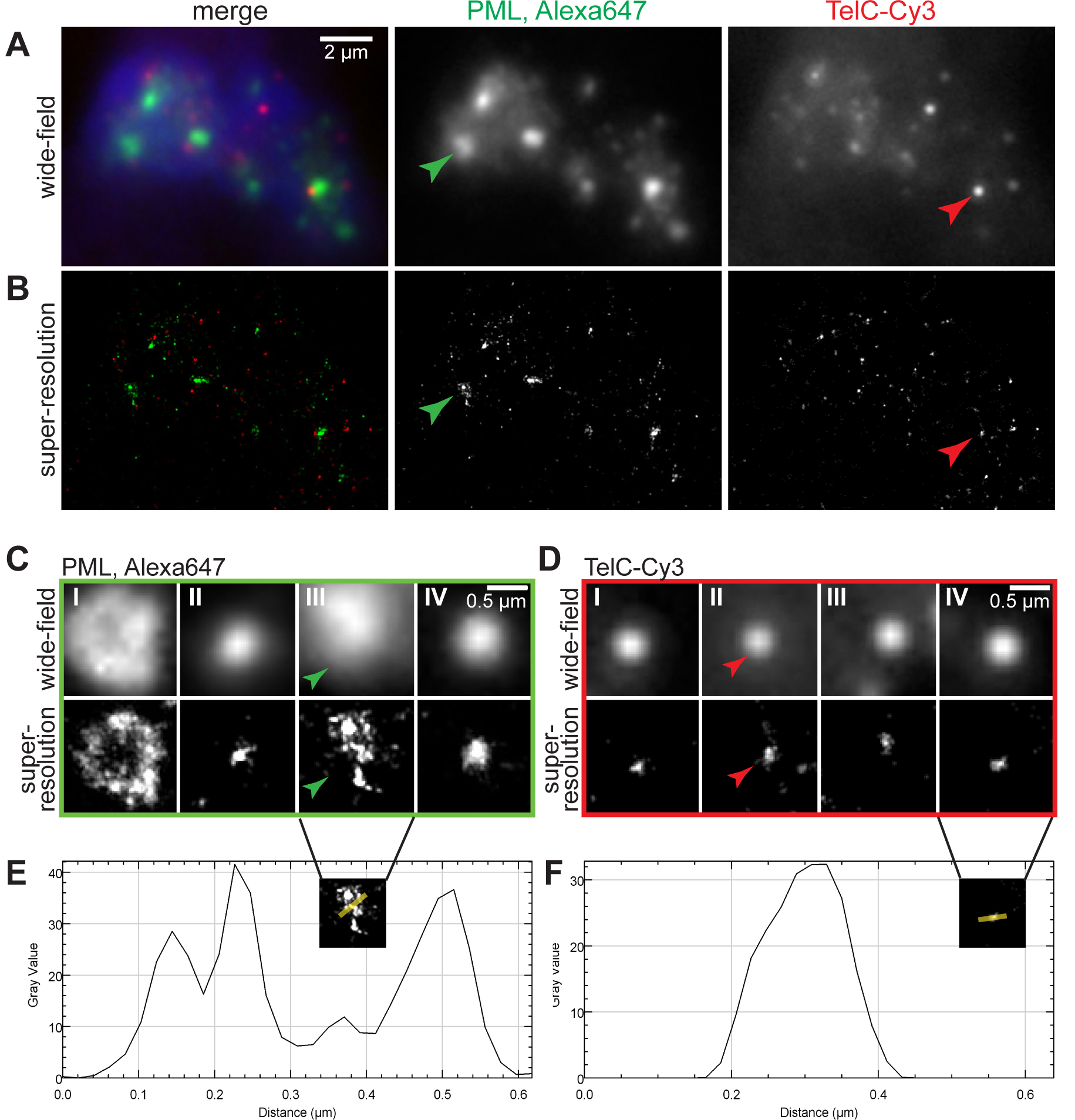
Super-resolution imaging of telomeres and PML-NBs in a pedGBM section. **A.** Wide-field overview image of PML-NBs (labeled with Alexa647) and telomeres (labeled with Cy3) in a FFPE pedGBM tissue section. **B.** Super-resolution reconstruction of the cell shown in A. **C.** Magnified regions of PML-NBs in different cells in the tissue section. In the top row, wide-field images are depicted. In the bottom row, the corresponding super-resolution reconstructions are shown. Green arrows mark a PML body identified in panels A and B. **D.** Magnified regions of telomeres in different cells in the tissue section (top row: wide-field images, bottom row: super-resolution reconstructions). Red arrows mark a telomere identified in panels A and B. **E.** Line plot of the PML body shown in panel C-III. **F.** Line plot of the telomere shown in panel D-IV. The plotting direction in E and F is indicated by the yellow line in the insert.

## 4. Concluding remarks

The identification of telomere features in primary tumor tissue sections is becoming increasingly relevant for patient stratification in several cancer types. Here, we have introduced an integrated workflow for automated 3D high-resolution image acquisition and analysis to dissect telomere features from tissue sections. Using these methods, telomere length distribution patterns in single pedGBM tissue sections as well as on a prostate cancer TMA could be determined. We show that the automated acquisition and image analysis of FISH-stained TMAs can be used to discriminate telomere length distributions in different patient samples. It is noted that telomere shortening in both prostate cancer cells and surrounding stromal cells have been suggested as prognostic markers in this disease that is associated with tumorigenesis [29, 47–49]. It was found that short or heterogeneous telomere lengths in tumor cells and short telomeres in cancer-associated stromal cells are correlated with a poorer prognosis and higher risk of cancer recurrence. Further, the simultaneous detection of PML protein and telomeres and the workflow proposed here enables the quantitative evaluation of APBs. These nuclear subcompartments are a hallmark for ALT positive tumor cells, and thus provides information about the active TMM. This is of particular interest for pedGBMs were recent studies indicate that the activation of ALT is linked to differences in clinical outcome and therapy response [22]. Thus, the correlation of the 3D-TIM based analysis of telomere features of tissue sections with clinical data provides an additional readout that can be integrated into patient stratification schemes. In addition, the application of super-resolution microscopy techniques to the analysis of cancer tissue sections is a novel and hardly exploited new possibility to investigate cellular substructures with respect to cancer-specific aberrations. With the fluorescent labeling scheme used here, a multi-mode and multi-scale imaging approach that combines wide-field microscopy, CLSM, and SMLM within a single instrument can be applied to primary tumor samples. By switching back and forth between the three different imaging modalities, imaging speed and resolution can be optimized for acquiring the specific datasets needed. We anticipate that this approach will also be useful for other applications that investigate the structural organization of healthy cells within their endogenous tissue environment.

## Acknowledgments

The work was funded by the German Federal Ministry of Education and Research (BMBF) within project CancerTelSys (grant number 01ZX1302) in the e:Med program. The ViroQuant-CellNetworks RNAi Screening Facility was supported by the CellNetworks-Cluster of Excellence (grant number EXC81).

